# Aberrant insula activity to negative and reduced learning from positive prediction errors as mechanisms underlying maladaptive self-belief formation in depression

**DOI:** 10.1101/2024.05.09.593087

**Authors:** Nora Czekalla, Alexander Schröder, Annalina V Mayer, Janine Stierand, David S Stolz, Tobias Kube, Christoph W. Korn, Ines Wilhelm-Groch, Jan Philipp Klein, Frieder M Paulus, Sören Krach, Laura Müller-Pinzler

## Abstract

Maladaptive self-beliefs are a core symptom of major depressive disorder. These beliefs are perpetuated by a negatively biased integration of self-related feedback. Understanding the neurocomputational mechanisms of biased belief updating may help to counteract maladaptive beliefs and the maintenance of depression. The present study uses a belief-updating task and functional neuroimaging to examine the neurocomputational mechanisms associated with self-related feedback processing in individuals with major depression and matched healthy controls. We hypothesized that increased symptom burden in depression is associated with negatively biased self-belief updating and altered neural tracking of social feedback. Our findings show that depression is related to reduced incorporation of unexpected positive feedback, with higher symptom burden alongside heightened insula reactivity to unexpected negative feedback. The interplay of increased neural responsiveness to negative feedback and the reduced learning from positive feedback provide new insights into cognitive distortions in depression and may explain the persistence of maladaptive self-beliefs and, thus, the maintenance of depression.

## Introduction

Maladaptive, mostly negatively biased, and highly rigid self-beliefs are prominent in various mental disorders^1^. Humans act on their beliefs as if they reflect reality^2^, which makes maladaptive beliefs a key maintaining factor of mental disorders^1^. These beliefs are a product of our learning history and are constantly updated in the face of incoming information that deviates from expectations^2–4^. Maladaptive beliefs in depression often center on a pessimistic view of oneself, like one’s ability or one’s future^1^. When the emphasis of this internal model is strong^5^, one becomes insensitive to the external context, such as contradictory positive feedback information^6^. To understand the computational mechanisms of maladaptive self-belief formation^7,8^, we assessed individuals with depression and matched healthy controls in a functional magnetic resonance imaging (fMRI) study. We used the Learning Of Own Performance (LOOP) task^7–9^ with trial-by-trial performance feedback to make participants form novel beliefs about their abilities. On their way to their new self-beliefs, we investigated the brain’s response to unexpected positive or negative performance feedback and measured accompanying affective states.

Negative self-beliefs are typical characteristics of depression and are maintained by negatively distorted information processing^10^. Accordingly, more negative belief-updating about one’s future, abilities, or social popularity has been observed in experimental settings in depression^11,12^. More recently, mental disorders have been conceptualized as an interplay of maladaptive prior internal models with a maladaptive response to prediction errors^5,13^, which is the mismatch between the expectation of any future event and the actual outcome. The persistence of negative beliefs in depression has been related to reduced updating after positive prediction errors^6^, which means higher insensitivity to the context when incoming information is better than expected. Cognitive immunization against this unexpected positive information is a cognitive mechanism that hinders the change of negative beliefs^6,14^. It reflects a reappraisal of unexpected feedback in a way that the initial expectation is maintained, for example, by devaluing positive feedback as an exception or by questioning its credibility.

The experience of prediction errors is linked to the experience of affect^15^. The current affective state can influence the updating of self-beliefs^8,16,17^, which, in turn, contributes to how the individual feels. This results in a recursive influence of affect and belief-updating on each other^16,17^. Accordingly, the experience of more negative and less positive affect during learning has been linked to negatively biased updating of self-beliefs^8,18^. Depression is characterized by predominantly negative affect^19^. Correspondingly, the negative learning history is particularly present in this pervasive mood disorder, which makes a negative interpretation of incoming information appear especially plausible^6^. This negative interpretation bias can impact the integration of incoming information and trigger particularly strong emotional responses^20^ in line with the negative interpretation, which further contributes to negative affect. When negative affect is experimentally induced, individuals show reduced flexibility in adjusting their beliefs in a positive direction^14^.

The entanglement of negatively biased belief updating and affective experiences also manifests on the neural systems level^8^. In the context of depression, most studies focused on the neural processing of reward prediction errors and behavioral adaptation following rewarding information unrelated to a self-belief^21^. In line with the persistence of negative beliefs against conflicting positive information, many studies reported reduced reward learning^22^ and reduced reward-related prediction error signaling in the ventral striatum and other brain regions, such as the ventral tegmental area, anterior cingulate cortex, and hippocampus in depression^23^. However, other studies found no support for altered reward learning^24,25^ or prediction error signaling in depression^25,26^. Similarly, in response to negative prediction errors, some studies reported increased activity^27^, while others did not^25^.

When processing self-related feedback, in particular, individuals with depression exhibit heightened sensitivity to negative feedback, such as social rejection^28^. When receiving negative feedback, they show increased activity in the amygdala, anterior cingulate cortex, left anterior insula, and left nucleus accumbens^28–30^. This heightened neural reactivity is associated with more negative self-perceptions, negatively biased information processing^29^, and lower self-esteem^28^. Altered insula activity in individuals with depression has been discussed as a neural correlate of more painful emotions^31^ and maladaptive emotion regulation in depression^29^. Additionally, inducing a depressed mood in healthy participants increases insula reactivity to negative information^32^. In individuals with social anxiety, which is also characterized by negative self-beliefs^33^ and often comorbid with depression, activity in the insula mediated the effect of negative social feedback on self-belief updates^34^. These findings highlight the crucial role of emotion-related brain regions in processing self-related feedback, setting the stage for examining how such neural responses contribute to the formation of maladaptive self-beliefs in depression.

While prediction error processing has been widely studied in depression using reinforcement learning tasks, much less is known about its role in the formation of self-beliefs^16^. The current study builds on previous work by applying a well-established computational approach of trial-by-trial self-belief formation, the LOOP task^7–9^, to a clinical sample with depression. To understand the mechanisms by which people arrive at their maladaptive self-beliefs, we aimed to test whether 1) depression is related to biased updating of self-beliefs about one’s ability in response to positive and negative prediction errors, and 2) whether processes of belief formation are underpinned by altered neural prediction error processing in depression. To investigate associations with prediction error processing in the brain, regions of interest (ROIs) in the insula and amygdala were defined based on our previous study with the same paradigm^8^. Based on other studies on reward learning and depression, we included the ventral striatum as an additional ROI. In line with negative self-beliefs and distorted information processing in depression, we expected more negatively biased self-belief formation accompanied by an unbalanced neural processing of positive and negative prediction errors. As we previously linked prediction error processing mapped in the insula and amygdala to biased self-belief formation and affect^8^, we expected depression-related alterations in these regions.

## Results

Participants diagnosed with major depressive disorder (MDD; *n*=35) and healthy control (CON) participants (*n*=32) completed the LOOP task^7,8^ in the MRI (for sample characteristics, see Supplementary Table 1a-c). In this task, participants estimated attributes of objects in two out of four different estimation categories (e.g., heights of buildings). On each trial, they received manipulated performance feedback relative to an alleged reference group (see Methods). The feedback referred to their own (Self) or another person’s estimation abilities (Other, Fig. 1A), who allegedly performed the task in an adjacent room. By linking estimation categories either with mostly “better than expected” or mostly “worse than expected” feedback, participants were led to form novel ability beliefs. The emotional response to self-related feedback was assessed with two ratings of happiness, pride, and embarrassment during the task. To describe the belief formation process, we modeled the changes in participants’ expected performance through updates from prediction errors for each participant. Consistent with our previous studies^7,8,35^, the winning model for both the depression and healthy control groups showed higher learning rates for negative compared to positive prediction errors (PE) in forming novel self-beliefs. This negativity bias was not present when participants formed beliefs about the other person’s ability (PE-Valence x Agent interaction: *t*(195)=-3.29, *p*<.001; Supplementary Table 2; negativity bias: post-hoc ɑ_Self/PE-_ vs ɑ_Self/PE+_; *t*(66)=4.78, *p*<.001; complete model space in Supplementary Table 3, for model selection, see Supplementary Note 1, Supplementary Table 4).

**Figure 1.**
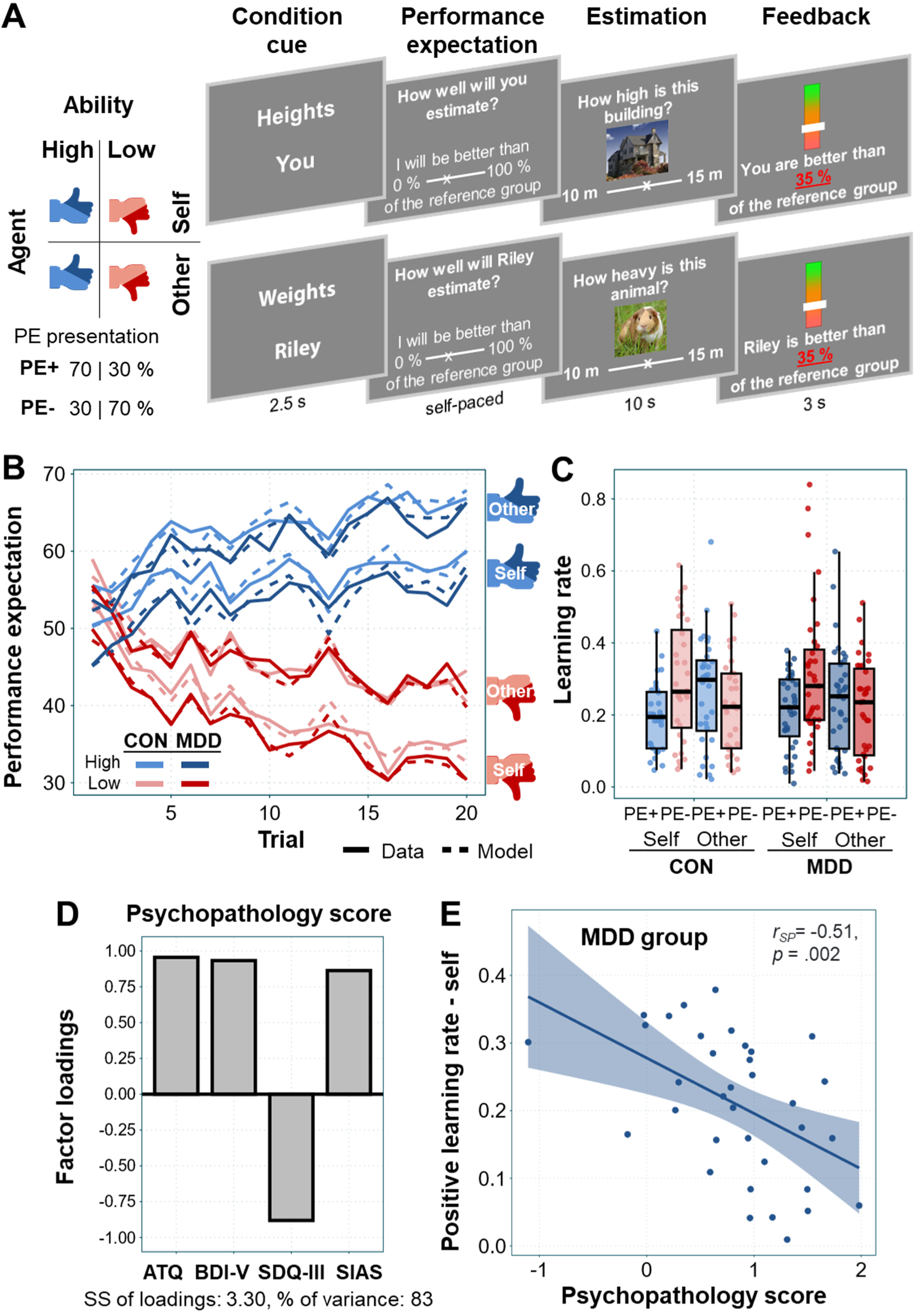
Trial sequence of the Learning-Of-Own-Performance (LOOP) task, experimental conditions, and behavioral results. **A** Low and High Ability conditions of alleged estimation performance for Self and Other (Agent condition) were presented in pseudo-randomized order. Feedback with controlled PEs: High Ability: 70 %, Low Ability: 30 % of the trials with planned positive PEs (the actual percentage of positive/ negative PEs could slightly differ, e.g., if the feedback had been out of range). Sequence of one self- and one other-trial from left to right: At the beginning of each trial, a screen displays the estimation category of the upcoming trial and whether participants have to perform a self-trial or an other-trial. This is followed by a rating of the expected performance (performance expectation). Next, participants are asked to estimate the specific attributes of objects (estimation), which is followed by a manipulated performance feedback (feedback). **B** Predicted and actual ratings of expected performance for both groups over time. The ratings indicate that participants update their expected performance (solid lines) according to the feedback, thus forming a belief about the performance levels in the different estimation categories. The winning model’s predicted values (dashed lines) with separate learning rates for Agent and PE valence captured the participants’ behavior, as indicated by a close match of actual ratings and predicted values. **C** Learning rates of the winning model show a bias towards increased updating in response to negative PEs (PE-) in contrast to positive PEs (PE+), specifically for self-beliefs in both groups. Colored bars indicate the first and third quartiles of the data; the line marks the median. Whiskers extend from the upper/ lower box borders to the largest/ smallest data point at most 1.5 times the interquartile range above/ below the respective border. MDD=Major Depressive Disorder (n=35), CON=control group (n=32). **D** Factor loadings of the principal component analysis, by which one factor, a summary measure of symptom burden, was extracted from the questionnaire variables (Automatic Thought Questionnaire, ATQ, Beck’s depression inventory, BDI-V, Self-Description Questionnaire-III, SDQ-III subscale scores, Social Interaction Anxiety scale, SIAS). SS=sum of squares. **E** Correlation of the positive learning rate (ɑ_Self/PE+_) and the Psychopathology score in the clinical sample. Spearman correlation coefficient, line for displaying purposes.

### No learning differences between individuals with depression and healthy controls

When comparing learning rates between the two groups, there was no difference in the overall extent of learning (main effect Group *t*(65)=-.30, *p=.*764) and no difference in the extent of the negativity bias (PE-Valence x Agent x Group interaction: *t*(195)=.18, *p=.*861; Fig. 1B and C, Supplementary Table 2). This is in line with the analysis of the expectation ratings in a model-agnostic approach (Supplementary Note 2, Supplementary Table 5). It suggests that people diagnosed with depression do not differ from healthy controls when forming novel beliefs in response to self-related performance feedback.

### Disregard of positive feedback with higher symptom burden

We additionally assessed self-reported symptom burden with a general measure of depressive symptoms (BDI^36^) and a specific measure of cognitive symptoms (ATQ^37^), as well as a measure of social anxiety (SIAS^38^) and self-esteem (SDQ-III^39^) as transdiagnostic markers, already shown in association with biased self-belief formation before^7^. Variability of symptoms was well explained by a single principal component (83% variance explained), reflecting general depressive and socially anxious psychopathology, akin to other recent approaches^40–42^ (see Fig. 1D and Methods section).

Within the clinical sample, symptom burden was associated with reduced self-belief updating following positive prediction errors (MDD: positive learning rate *r*ɑ_Self/PE+_=-.51, *p*=.002; for comparison, negative learning rate *r*ɑ_Self/PE-_=.03; difference of absolute values: *z*=2.1, *p=*.035; Fig. 1E, Supplementary Fig. 1. For more detailed correlations with symptom scores, see Supplementary Fig. 2). This indicates that individuals with higher symptom burden integrate positive feedback less, while learning from negative feedback remains unaffected.

### Imbalanced prediction error signaling in depression during feedback processing

Next, we examined the neural processing of self-related positive and negative PEs as a basis of self-belief formation (continuous PE effect). When comparing individuals with depression and healthy controls, a significant Group x PE-Valence interaction in the right anterior and posterior insula indicated an imbalance of negative relative to positive prediction error signaling in the clinical sample but not in healthy controls (anterior: *p*=.045, posterior: *p*=.030, family-wise [FWE] corrected at peak level within ROIs, Fig. 2, Supplementary Table 6, for baseline activations of PE effects, see Supplementary Table 7). This effect was not present in the amygdala and ventral striatum ROIs. There was no group difference for neural activity with more positive self-related prediction errors. For negative prediction errors, post-hoc t-tests suggest a stronger activity for individuals with depression than healthy controls (two-sample t-tests within ROIs, anterior: *p*=.059, posterior *p*=.014). The findings suggest that individuals with depression are more sensitive to negative prediction errors at the neural level (for general feedback effects of Self/Other, PE+/PE-feedback categories, see Supplementary Fig. 3, for whole brain analysis of negative compared to positive PE processing [continuous PE effect], see Supplementary Fig. 4).

**Figure 2.**
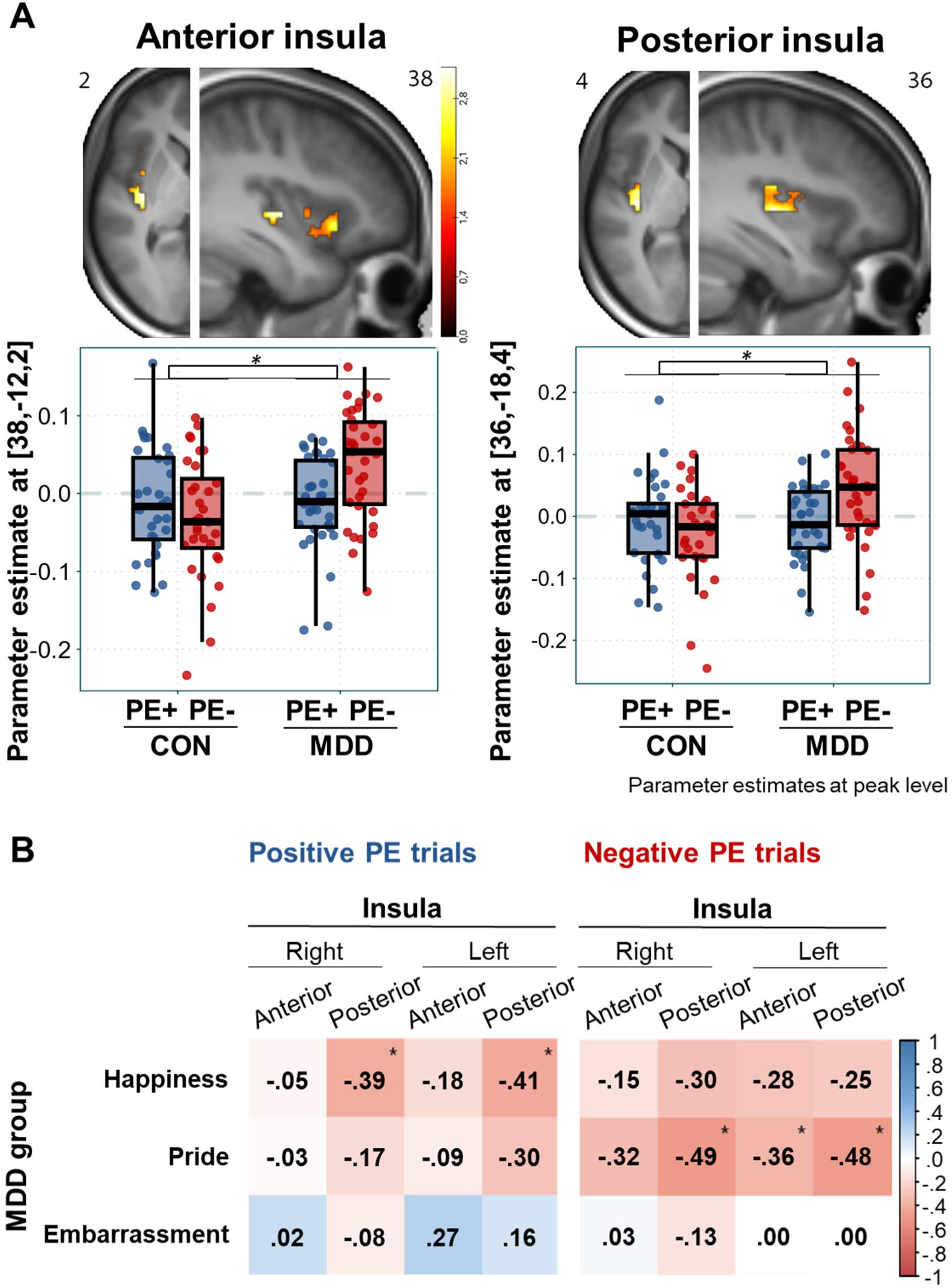
Differential insula tracking of PE valence and correlations with affective state. **A** The interaction (MDD[PE-]>MDD [PE+])>(CON [PE-]>CON [PE-]) shows a greater difference between PE- vs. PE+ in the MDD compared to the CON group (FWE corrected within ROIs at peak level; ROI selection guided by previous fMRI study^8^). CON=control group (n=32), MDD=Major Depressive Disorder (n=35). The plot depicts the parameter estimates of the Group [CON, MDD] x PE-Valence [PE+, PE-] interaction at peak level for the two regions of interest in the right insula. Brain plot: uncorrected *p<*0.05 within ROI for display purposes; see Supplementary Table 6 for FWE corrected statistics. **B** Spearman correlation between the affect ratings and brain activity within insula ROIs in response to positive and negative prediction errors (categorical PE effect) in the clinical sample. * *p* < .05.

Subsequent exploratory analysis within the clinical sample suggests that activation in these same regions is associated with reduced positive affect experienced during the task (for affect ratings in both groups, see Supplementary Table 8). In negative prediction error trials, stronger activation corresponds to lower pride (Spearman correlation between averaged parameter estimates within insula ROIs, right-anterior: *r*=-.32 [*p*=.058], posterior: *r*=-.49 [*p*=.003], left-anterior: *r*=-.36 [*p*=.034], posterior: *r*=-.48 [*p*=.004], Fig. 2, Supplementary Fig. 5). Correlations with happiness show a similar negative trend but do not reach significance. Notably, for positive prediction errors, stronger activation also aligns with decreased happiness, following the same pattern (right-posterior: *r*=-.39 [*p*=.019], left-posterior: *r*=-.41 [*p*=.014]). This pattern may suggest that deviations in either direction are processed negatively. Correlations with symptom burden within the clinical sample show the same directional pattern as the group comparison, indicating that higher symptom burden is associated with greater activation following negative prediction errors, although not statistically significant (negative PE trials: insula ROIs *r*=.23-.31 [*p*=.07-.18]; no meaningful correlations for positive PE trials: *r*=-.01-.03 [*p*=.79-.97]). Generally, while participants receive recurrent performance feedback, insula activation in negative prediction error trials is correlated with the learning behavior as indicated by the negative learning rates (see Supplementary Fig. 6).

Together, these behavioral and neural findings indicate that individuals with depression show impaired integration of positive feedback, paralleled by heightened neural sensitivity to negative feedback, highlighting a valence-specific imbalance in self-related learning mechanisms.

## Discussion

Maladaptive self-beliefs are a core feature of depression, yet the neurocomputational mechanisms underlying their formation and persistence remain unclear. Here, we examined the role of prediction error processing during belief formation using a computational modeling approach. Our findings suggest that individuals with depression exhibit reduced incorporation of positive self-related prediction errors with higher symptom burden alongside heightened insula reactivity to negative self-related prediction errors, potentially reinforcing maladaptive self-beliefs. These results provide new insights into cognitive distortions in depression and the neural mechanisms sustaining negative self-perceptions.

Both, the depression and the healthy control group show a self-specific negativity bias in belief updating. Our finding aligns with recent reward-learning studies of other groups that also do not find different learning patterns in depression and healthy controls^24,25^ and extends it to the field of self-related learning. This observation challenges the notion of fundamental differences in basic learning mechanisms^43^. A key factor in understanding the discrepancy between earlier studies that specifically target self-beliefs and do find group differences, and our study, may be the task’s unique design. Our task demands active task performance, creating the perception of control over feedback. Such task engagement can elicit specific self-related motivations, such as the motivation to improve one’s ability and receive positive feedback^44^ or the motivation to avoid failure^45^. Fear of failure can shift the focus toward negative feedback as a form of threat monitoring^46^. Additionally, we selected a domain with little prior experience to encourage novel belief formation, potentially leading to low confidence and a general sense of uncertainty during task performance. We consistently observed a negativity bias in belief updating, also in healthy individuals. It suggests that the transition from a healthy level of negative to a maladaptive self-belief formation may be subtle, becoming more evident in dimensional analyses rather than categorical group comparisons.

In the clinical sample, self-belief updates following positive prediction errors were reduced with increased symptom burden. One explanation is that positive feedback simply is less salient and therefore might draw fewer attention^47^. Alternatively, cognitive mechanisms could actively contribute to the devaluation of positive feedback, resulting in a cognitive immunization against it^48^. These findings suggest that sensitivity to context information, such as social feedback, diminishes as symptom burden increases, potentially due to an overemphasis on an entrenched negative internal model. As a result, learning from positive prediction errors is reduced, and beliefs become biased toward prior models in a confirmatory manner^49^.

On the neural system level, we observed stronger tracking of negative prediction errors in the insula in the clinical sample, despite no group differences in behavior. The stronger neural response to negative prediction errors aligns with a general neural sensitivity to negative (social) prediction errors in the insula^8^, as well as heightened insula reactivity in depression^50^ and social anxiety^34,51^. Given that insula reactivity to negative prediction errors was linked to reduced positive affect during task performance, the observed group difference may reflect heightened emotional responsivity in depression. This corresponds to the overall reduced positive affect in individuals with depression^19^ and underscores the notion of affected beliefs^8^ - the entanglement of beliefs and affect.

The stronger neural and possibly affective reaction to more negative self-related feedback can also be discussed in the context of emotional schemas and their role in shaping emotional responses. Emotional schemas are formed in highly arousing emotional events and can be triggered by learned signs associated with the events. This results in automatic and exceptionally intensive emotional responses^20^. Depression is more likely related to early maladaptive schemas rooted in an aversive learning history, such as experienced social devaluation in response to failure^52^. Therefore, individuals may have emotional schemas that are particularly sensitive to negative self-related feedback, which results in a stronger emotional response and, thus, an overall lower positive affect during task performance. In turn, the stronger reaction could indicate that the corresponding negative learning history is particularly present, so that the emphasis on a negative internal model is strong, making new information that deviates in the positive direction seem more implausible. The reduced positive updating found on the behavioral level may support this notion, though there was no such effect for positive prediction errors in the brain. However, the brain’s response is measured when the feedback is given, while the updating of the belief likely occurs at a later time, somewhere between feedback, feedback processing, interpretation, and indication of the new expectation. Given the alternating order of the positive and negative ability conditions, belief updates measured in behavior may represent an amalgam of several trials. If the reaction to negative feedback in the brain is strong, cognitive appraisal mechanisms might come into play and influence self-belief updating even following positive prediction errors. To better understand task-specific negative affective reactions to self-related feedback, trial-by-trial fluctuations in emotions should be measured in future studies.

Our findings of unaltered positive prediction error processing in the VS align with some prior research in depression, which has reported intact VS activity during reward learning tasks^25,26^. Coupled with the above-mentioned similar learning behavior in both groups, our results challenge the hypothesis of altered VS reactivity and prediction error learning deficits in depression^22,53^. Instead, the findings suggest that the neural processing of positive self-related prediction errors as a basis for forming positive self-beliefs remains intact in depression, at least at this initial stage of processing.

What can be learned from the current findings for the therapeutic practice? The findings can offer insights into addressing maladaptive beliefs therapeutically. As several aspects are already present in existing psychotherapy practices, our findings offer new perspectives into bridging the gap between research results and therapeutic interventions. Rather than directly challenging the negative self-concepts of patients with depression (e.g., by cognitive restructuring), they can be supported in becoming more context-sensitive so that positive feedback that contradicts the internal model can be incorporated. This might be done by practicing skills that help to become less emotionally reactive to established negative self-beliefs, e.g., with the idea of defusion from Acceptance and Commitment Therapy (ACT^54^), the detached mindfulness concept from Metacognitive Therapy^55^, or increasing sensitivity to the present context, as e.g., being implemented in mindfulness skills in ACT or mindfulness-based cognitive therapy^56^. To update a maladaptive belief, it may be necessary for a therapist to thoroughly examine the emotional and cognitive meaning attributed to new experiences, support the understanding and regulation of intense maladaptive emotional reactions, and identify potential cognitive distortions, such as retrospectively devaluing positive experiences. These mechanisms can be discussed with patients as a factor that maintains and continuously corroborates a negative self-image.

In conclusion, this study provides insight into neurocomputational mechanisms that contribute to maladaptive self-belief formation in depression. Our findings highlight a heightened neural response to more negative self-related feedback in the insula, potentially reflecting a more affectively charged processing style. Alongside reduced consideration of positive feedback with increasing symptom burden, it may promote negatively biased self-belief formation in depression in the long run. In the absence of fundamental deficits in learning behavior in depression, this effect underscores the importance of accounting for within-group variance and symptom burden within a clinically relevant range. Insights from the interaction between cognitive processes, affective experiences, and symptom burden towards forming maladaptive self-beliefs can be of great importance for future therapeutic intervention strategies.

## Methods

### Participants

The study was approved by the ethics committee of the University of Lübeck (AZ 18-066), was carried out in accordance with the ethical guidelines of the American Psychological Association, and all participants gave written informed consent. All participants were adults above the age of 18 and fluent in German. The clinical sample (*n*=35, 9 females, aged 20–55 years; *M*=34.23; *SD*=10.54) was recruited from the Department of Psychiatry and Psychotherapy at the university clinic Lübeck. The goal in recruiting was to obtain a naturalistic sample with variance in depressive and social anxiety symptoms. Inclusion criteria were a diagnosis of depression (F32.1/2, F33.1/2; given by medical or psychological therapists at the clinic); comorbid symptoms of social anxiety (F40.1, F60.6) were preferentially included but were not a necessary inclusion criterion. Exclusion criteria were acute suicidality, an acute psychotic state, schizophrenia, schizotypal disorder or delusional disorder, bipolar disorder, primary diagnosis of substance abuse or substance dependence, narcissistic, histrionic, or borderline personality disorder, and neurological disease. Subjects of the control group (*n*=32, 10 females, aged 18–55 years; *M*=34.25; *SD*=10.23, no group differences in age or gender distribution *p*>.8, Supplementary Table 1a) were recruited at the university campus of Lübeck or via public notices. The exclusion criterion was mental illness or neurological disease. Individuals interested in the study were screened for symptoms of depression through a telephone interview, and the second part included a general screening for other mental illnesses adapted from a structured interview (SCID; for additional sample descriptions, see Supplementary Note 3, Supplementary Table 1).

### Learning Of Own Performance (LOOP) task

The LOOP task assesses how participants learn about their own or another person’s purported estimation ability through trial-by-trial performance feedback. It was covered as a cognitive estimation experiment. The task was previously validated in behavioral and fMRI studies on healthy participants^7,8,35^. Two participants were invited, one performing the task in the scanner and the other completing a behavioral version (data not included). When a second participant was unavailable, a confederate was introduced as the behavioral participant. Participants alternated between two trial types: actively performing the estimation task (*Self*-trial) or observing another person’s performance (*Other*-trial). The task involved estimating attributes in four categories: house height, animal weight, vehicle distance, and food quantity. Feedback was manipulated such that, for each agent, one category was paired with predominantly positive feedback (*High Ability*), while the other was paired with mostly negative feedback (*Low Ability*), resulting in four feedback conditions: *Self-High*, *Self-Low*, *Other-High*, and *Other-Low* (20 trials each). Each feedback condition was linked to a specific estimation category. Trials of all conditions were intermixed in a fixed order, preventing more than two consecutive trials of the same condition. Feedback, presented as percentiles relative to an alleged reference group of 350 former participants, was determined by a sequence of predefined PEs based on participants’ evolving ability belief. This was calculated as a moving average of the last five performance expectations per category, initialized at 50%. This approach ensured that PEs remained largely independent of performance expectations and that negative and positive PEs were relatively equally distributed across groups and conditions (see Supplementary Note 4).

Each trial began with a cue indicating the estimation category and agent. Participants then rated their performance expectation (in percentiles) and were incentivized with up to 6 cents per trial for accurate predictions of their performance. The estimation question was presented for 10 s, with responses made on a continuous scale within a predefined plausible range for each question. In *Other*-trials, participants also provided estimates to ensure comparable mental effort across both agent conditions. Subsequently, feedback was presented for 3 seconds (Fig. 1A). In *Other*-trials, the other’s feedback was visible, while their performance expectations and answers to the estimation questions remained private. Jittered inter-stimulus-intervals were presented following the cue (mean: 4*TR [0.992 s], range: 2–6*TR), estimation (mean: 4.5*TR, range: 2.5–6.5*TR), and feedback phase (mean: 6*TR, range: 4–8*TR) with jitters distributed in a uniform distribution with steps of 0.5*TR.

To assess affective responses, participants rated their current levels of embarrassment, pride, happiness, arousal, and tiredness on a continuous scale (0-100) four times, each following one trial per feedback condition. The two ratings following *Self*-trials were averaged to measure affective responses to self-related feedback. The task was implemented using MATLAB (Release 2015b, The MathWorks, Inc.) and the Psychophysics Toolbox (Brainard, D. H. [1997]. The Psychophysics Toolbox. *Spatial Vision*, *10*, 433–436). The task comprised two ∼25-minute sessions with a short break in between.

### Statistical analysis

#### Behavioral data analysis and modeling

##### Model-agnostic analysis

First, we checked for group differences in prior estimation ability beliefs (Supplementary Note 5, Supplementary Table 1b), other baseline characteristics like estimation experience and importance, and social perceptions of the other (Supplementary Table 1c). Then, we ran a model-agnostic analysis on the participants’ expected performance ratings for each trial. We employed a linear mixed model with a maximum likelihood estimation. The model included the factors Ability (High/Low) x Agent (Self/Other) x Group (MDD/CON) and Trial (20 trials) as a continuous predictor. Intercept, Ability, Agent, Trial, and Group were modeled as fixed effects; the intercept was additionally modeled as a random effect (Supplementary Note 2, Supplementary Table 5).

##### Computational modeling

Dynamic changes in performance expectation ratings were modeled using PE delta-rule update equations (adapted Rescorla–Wagner model^57^). The model space and the procedure of model fitting and selection have been implemented and further developed in our previous studies^7,8^. The model space included three main approaches: the Unity Model (single learning rate), the Valence Model (distinct learning rates for positive vs. negative prediction errors), and the Ability Model (distinct learning rates for ability conditions). A Mean Model served as a baseline, assuming stable values. Learning rates were estimated either separately for Self vs. Other or across both agents (for detailed model space description, see Supplementary Note 6, Supplementary Table 3). The winning model was a Valence Model with separate learning rates for Self vs. Other. It was further extended by adding a weighting factor that reduced the learning rates when feedback values approached the extremes of the scale (percentiles close to 0% or 100%), assuming that participants would perceive extreme feedback to be less likely than average feedback^58^. Since many variables encountered in everyday life approximately follow a normal distribution where extreme values are less probable, we assigned the relative probability density of the normal distribution to each feedback percentile value. A weighting factor *w* was fitted for each individual, indicating how strongly the relative probability density reduced the learning rates for feedback further away from the mean. The Weighted Valence Model was the winning in our last study^8^. The normal decay (ND) weighted by the weighting factor *w* was introduced in the learning models in the following way: (EXP=Performance expectation rating, FB=feedback, PE=prediction error, α=learning rate):

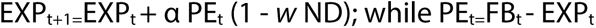

The initial beliefs about the own and the other participant’s performance (EXP_1_) were estimated as free parameters separately for Self and Other in both Ability conditions, resulting in four additional model parameters.

For model fitting, we used RStan, employing Markov Chain Monte Carlo sampling to estimate individual learning parameters. Across three chains, 2400 samples were drawn after 1000 burn-in samples (thinned by 3). Convergence was assessed via R^ values^59^, with effective sample sizes (*n_eff_*) > 1500 for most parameters and subjects. Posterior distributions were summarized by their mean for group-level analysis.

For model selection, we used leave-one-out cross-validation (LOO^60^) with Pareto smoothed importance sampling (PSIS) to assess the prediction accuracy of fitted models. K^ values evaluated PSIS-LOO reliability, showing only a few trials with insufficient values and, thus, potentially unreliable PSIS-LOO scores (Supplementary Table 4). Bayesian Model Selection on PSIS-LOO scores was performed on the group level for the whole sample and both sub-samples^61^ (Supplementary Note 1). The winning model was validated by replicating the model-agnostic analyses with its predicted data (see Supplementary Note 7; for model parameter correlations, see Supplementary Fig. 7).

##### Capturing symptom burden

For a dimensional perspective on depression, we assessed the severity of depressive symptoms with the BDI and ATQ. To address symptom burden in a broader context, we additionally assessed symptoms of social anxiety with the SIAS and self-esteem with the SDQ-III (subscale scores) as we could show a relationship with belief updating behavior and these two concepts in a previous study^7^. To reduce dimensionality, we ran a principal component analysis on our questionnaire data using the *R* package ‘*psych*’ (varimax rotation, component scores based upon the structure matrix [default], William Revelle (2023). psych: Procedures for Psychological, Psychometric, and Personality Research. Northwestern University, Evanston, Illinois. R package version 2.3.3, https://CRAN.R-project.org/package=psych). We extracted one component as a combined measure of depressive and socially anxious psychopathology (Fig. 1E).

##### Statistical analyses of learning parameters

To test whether the updating of self-beliefs was different between groups, learning rates of the winning models for positive and negative PEs (factor PE-Valence) and for Self and Other (factor Agent) were compared between the two groups in a PE-Valence x Agent x Group linear mixed model. For a more detailed understanding of the relationship between biased belief updating and symptom burden, we additionally performed Spearman correlations between the Psychopathology score and the self-related learning rates ɑ_Self/PE+_ and ɑ_Self/PE-_ within the clinical samples. Since we had a particular interest in the relationship between the Psychopathology score and the self-related positive learning rate as a potential measure of cognitive immunization against positive feedback, we additionally tested the absolute correlation parameter of ɑ_Self/PE+_ and Psychopathology against the one with ɑ_Self/PE-_ within the clinical sample (*r* package *‘cocor’* that implemented to correlation comparison approach by Pearson and Filon, 1898; Diedenhofen, B. & Musch, J. (2015). cocor: A Comprehensive Solution for the Statistical Comparison of Correlations. *PLoS ONE, 10*(4): e0121945). Statistical tests were performed two-sided. Statistical analyses on the behavioral data were performed using *R* (R Core Team [2022]. R: A language and environment for statistical computing. R Foundation for Statistical Computing, Vienna, Austria. https://www.R-project.org/).

##### FMRI data acquisition

FMRI data were collected at the Center of Brain, Behavior, and Metabolism at the University of Lübeck, Germany, using a 3 T Siemens MAGNETOM Skyra scanner (Siemens, München, Germany) with 60 near-axial slices. 1516 functional volumes (min=1400, max=1724) were acquired on average per session using echo planar imaging (EPI, TR=0.992 s, TE=28 ms, flip angle=60°, voxel size=3×3×3mm^3^, simultaneous multi-slice factor 4). A high-resolution anatomical T1 image was obtained for normalization purposes (voxel size =1×1×1mm^3^, 192 × 320 × 320 mm^3^ field of view, TR=2.300 s, TE=2.94 ms, TI=900 ms; flip angle=9°; GRAPPA factor 2; acquisition time 6.55 min).

##### FMRI data analysis

FMRI data were preprocessed and analyzed with Statistical Parametric Mapping 12 (SPM12, Wellcome Trust Centre for Neuroimaging, University College London). Field maps were recorded to obtain voxel displacement maps (VDMs) to correct for geometric distortions. EPIs were slice-time corrected, motion-corrected, and unwarped using the corresponding VDMs, co-registered with the T1 image, and normalized using the forward deformation fields as obtained from the unified segmentation of the anatomical T1 image. The normalized volumes were resliced with a 2×2×2 mm^3^ voxel size and smoothed using an 8mm, full-width-at-half-maximum isotropic Gaussian kernel. Functional images were high-pass filtered at 1/384 to remove low-frequency drifts.

A two-level, mixed-effects procedure was implemented for the statistical analyses. On the first level, a fixed-effects GLM included four regressors for the cue conditions (Ability: High vs. Low × Agent: Self vs. Other), weighted with the performance expectation ratings per trial as parametric modulator, four regressors for the feedback conditions (Agent [Self /Other] × PE-Valence [PE+/PE-], categorical PE effect), weighted with PE magnitude per trials (continuous effect of the unsigned PE values for each feedback condition), one regressor for the performance expectation rating phase, two for the estimation period for Self and Other, and one for the emotion rating phase. Parametric modulators were not orthogonalized; thus, each only explained their specific variance. Regressors were modeled with the duration as presented during the experiment (cue phase: 2.5s, performance expectation rating: individual reaction times with *M*=4.0s, *SD*=2,4, estimation phase: 10s, feedback phase: 3s, emotion rating phase: *M*=26.2s, *SD*=10.1). Six additional regressors were included to correct for head movement, and one regressor was included with a constant term for each of the two sessions.

On the second level, we first compared the brain activity for self- vs. other-related positive and negative PEs in the feedback phase (categorical PE effect) in a flexible factorial design (regressor for the four conditions Agent [Self /Other] × PE-Valence [PE+/PE-]) to test whether results of our previous study^8^ replicate (Supplementary Fig. 4). To compare the tracking of positive and negative self-related PEs captured by the parametric modulator of PE magnitude (continuous effect) between the MDD and CON group, we tested the PE-Valence × Group interaction in a flexible factorial design within our ROIs. We compared activity between MDD and CON separately for positive and negative PEs in post-hoc two-sample t-tests. In a subsequent exploratory analysis, we correlated brain activity in self-related positive and negative PE trials (categorical PE effect, extracted and averaged parameter estimates of all voxels within one ROI) with ɑ_Self/PE+_ and ɑ_Self/PE-_, respectively, as well as the Psychopathology score and the emotion ratings using Spearman correlation.

The selection of regions of interest was guided by our previous fMRI study with a healthy sample using the same paradigm^8^. As the insula has emerged as particularly relevant in this study, it was again defined as an ROI. We used unilateral insula ROIs as described in the three-cluster solution of Kelly and colleagues^62^, with their dorsal and ventral part combined into an anterior ROI and a posterior part. Other ROIs motivated by this study are in the amygdala (two unilateral ROIs from the AAL atlas definition in the WFU PickAtlas^63^). Since altered PE processing in the ventral striatum has been discussed a lot in the context of depression, a functional ROI within the ventral striatum (according to the SPM anatomy toolbox) capturing the processing of PE valence was additionally defined as ROI. It derived from a family-wise error *p<*.05 corrected baseline effect of the continuous signed self-related PEs of our previous study^8^.

FMRI results were family-wise error (FWE) corrected at peak level for the whole brain or within the ROIs. Coordinates are reported in the MNI space. Anatomical labels of all resulting clusters were derived from the SPM Anatomy toolbox, version 3.0 (Eickhoff, S. B., Stephan, K. E., Mohlberg, H., Grefkes, C., Fink, G. R., Amunts, K., & Zilles, K. [2005]. A new SPM toolbox for combining probabilistic cytoarchitectonic maps and functional imaging data. Neuroimage, 25(4), 1325-1335).

## Data and Code availability

The behavioral data and code used to generate the behavioral analyses are available at https://osf.io/gwkp3/ (Identifier: DOI 10.17605/OSF.IO/GWKP3). For a quick overview, see *R Markdown/Quarto* output document. Neural data and analysis scripts, as well as *R Stan* scripts to estimate the computational models are available from the corresponding author upon reasonable request.

## Acknowledgments

We are grateful to Rebecca Rocksien, Katharina Ohm, Ann-Kristin Schmidt, Jovana Lehmann-Grube, Alica Steinert, Leonie Christine Pasquier, and Yana Schwarze for their help with data collection. We thank Yana Schwarze for providing a Matlab analysis script to check the MRI motion parameters. The research was funded by the German Research Foundation (Temporary Positions for Principal Investigators: MU 4373/1-1; Sachbeihilfe KR 3803/11-1; 3803/14-1) and the Medical Department of the University of Lübeck (J21-2018).

## Author Contributions

L. M.-P., S. K., F. M. P., N. C. conceived and designed the experiments. N. C., A. S., J. S. collected the data, with the consultation of J. P. K. to recruit the clinical sample. N. C. analyzed the data under the supervision of L. M.-P., with the consultation of F. M. P.. N. C., A. S., A. V. M., T. K., F. M. P., S. K., and L. M.-P. discussed the data analyses and interpretation of the results. N. C. wrote the original draft of the manuscript. L. M.-P. and S.K. supervised and edited the paper. A. S., A. V. M., J. S., D. S. S., T. K., C.W.K., I. W.-G., J. P. K., F. M. P. reviewed and edited the paper.

