## Supplementary_Information for "Aberrant insula activity to negative and reduced learning from positive prediction errors as mechanisms underlying maladaptive self-belief formation in depression"

**Czekalla et al.**

### Supplementary Notes

**Supplementary Note 1: Model selection.**

The weighted Valence Model (M6), including separate learning rates for positive and negative prediction errors for Self vs. Other, received the highest sum PSIS-LOO score (approximate leave-one-out cross-validation [LOO] using Pareto smoothed importance sampling [PSIS][1]) out of all models (for the structure of the model space see Supplementary Table 3, for all PSIS-LOO scores, see Supplementary Table 4). In addition, Bayesian model selection[2] in the whole sample and the clinical and control sub-samples resulted in a protected exceedance probability of $pxp > .999$ for this model and a Bayesian Omnibus Risk of $BOR <$ .001. The expected model frequency was 55.07 (MDD: 27.51, CON: 27.48). Thus, the Weighted Valence Model was selected for all further analyses of learning parameters, allowing for a comparison of valence-specific learning rates (for parameter correlations, see Supplementary Fig. 7).

**Supplementary Note 2: Model-agnostic behavioral analyses.**

We performed a model-agnostic analysis on our performance expectation ratings over time to capture the basic effects of belief updating in our observed data and compare them between the two groups. The Trial x Ability condition x Agent condition x Group linear mixed model showed a significant main effect of Ability condition ($t(5279) = 4.4, p < .001$) and interaction of Trial x Ability condition ($t(55279) = 10.3, p < .001$), indicating that participants adapted their performance expectation ratings according to the presented feedback in each Ability condition (see Fig. 1B). Moreover, there was a significant main effect of Agent ($t(5279) = -2.9, p = .004$) and a significant interaction of Trial x Agent ($t(5279) = -2.6, p = .010$), indicating that participants evaluated their performance increasingly more negatively over time than the other's performance. The main effect of Group and all interactions which included Group were not significant (for a full results table, see Supplementary Table 5).

**Supplementary Note 3: Additional sample description.**

In the clinical sample, 28 participants were outpatients, and seven were inpatients. Twenty-seven of the outpatients were in a day treatment program for depression (8 weeks, weekday mornings to afternoons) with multi-professional therapy. All patients received psychotherapy, and $n = 28$ received additional pharmacotherapy. Measurement day was, on average, 2.4 weeks ($SD = 2.0$) after admission to the clinic.

We initially recruited 85 participants in total but had to exclude 18 participants. In the clinical sample, we excluded five due to technical problems, four due to premature termination of the study, and two participants who did not attentively complete the task until the end (e.g., ratings indicated that they stopped responding). In the control group, we excluded three participants who did not believe the cover story of the task and/ or did not attentively complete the task; one was excluded due to technical problems, one due to a premature termination of the study, and two due to clinically relevant BDI scores (> 35), despite screening before.

**Supplementary Note 4: Means and frequencies of presented prediction errors in both groups.**

As outlined in the methods section, feedback was determined using a series of predefined prediction errors (PEs) based on the participants' evolving ability beliefs. These beliefs were estimated as a moving average of the five most recent performance expectations within each category. In contrast, the actual PE was computed from the current performance expectation rating on a given trial, which could lead to slight deviations between the predefined and the actual PE distributions. In addition, feedback values falling outside the permissible range were replaced.

In the clinical sample, the PEs were distributed as follows: In the Self condition, mean positive PE = 14.2, *SD* = 1.5 (mean frequency = 20.4); mean negative PE = −12.8, *SD* = 1.8 (mean frequency = 19.1); and in the Other condition, mean positive PE = 13.6, *SD* = 1.3 (mean frequency = 19.0); mean negative PE = −14.0, *SD* = 1.8 (mean frequency = 20.3). In the control group: Self condition, mean positive PE = 13.9, *SD* = 1.4 (mean frequency = 20.2); mean negative PE = −13.2, *SD* = 1.4 (mean frequency = 19.3); Other condition, mean positive PE = 13.8, *SD* = 1.7 (mean frequency = 18.7); mean negative PE = −13.5, *SD* = 1.2 (mean frequency = 20.7). There were no group differences in mean PE for all four PE conditions, all $p > 0.2$).

**Supplementary Note 5. Group differences in global but not in specific prior beliefs.**

Before comparing the process of belief formation between the two groups, we checked if there were already group differences in participants' prior beliefs regarding their estimation abilities. While individuals with depression showed a lower evaluation of the own abilities in general (*SDQ-III*, $t(45) = 8.23$, $p < .001$) as well as their general estimation abilities before the task ($t(62) = 2.1$, $p = .04$). However, when asking more specifically for the performance in a certain estimation category or even more specific for the performance expectation in the upcoming first trail there were no group difference, neither for self (category-specific estimation ability: $t(64) = 1.04$, $p = .301$; 1. performance expectation rating: $t(56) = 1.83$, $p = .072$) nor for other-related prior beliefs (category-specific estimation ability: $t(64) = 0.82$, $p = .415$; 1. performance expectation rating: $t(59) = 1.39$, $p = .17$). This allows for a comparison of the learning process between groups independently of the prior beliefs (for a complete table of prior beliefs with group comparisons see Supplementary Table 1b).

**Supplementary Note 6. Description of the model space.**

The model space consisted of three main models, each with different assumptions regarding biased updating behavior when forming novel beliefs. The simplest learning model (Unity Model) employed a single learning rate for all conditions for each participant, thus assuming no learning biases. The Valence Model included separate learning rates for positive and negative prediction errors across both ability conditions, suggesting that the valence (positive vs. negative) of prediction errors influences belief formation. The Ability Model incorporated distinct learning rates for each ability condition, indicating context-specific learning. To compare whether the participants' performance expectation ratings can be better explained in terms of prediction error learning compared to assuming stable values within each ability condition, we included a simple Mean Model with a mean value for each task condition (for model space, see Supplementary Table 3).

**Supplementary Note 7: Posterior predictive checks - Behavioral analyses on the predicted data.**

We repeated the analyses (Supplementary Note 2) using the predicted data from the winning model to see whether our winning model captured the core effects in our model-agnostic analysis. We could reproduce all effects from the with the predicted data (main effect Ability condition: $t(5279) = 4.04$, $p < .001$, main effect Agent condition: $t(5279) = -4.09$, $p < .001$, interaction Trial x Ability condition: $t(5279) = 14.2$, $p < .001$, interaction Trial x Agent condition $t(5279) = -2.51$, $p = 0.012$). This confirms that the winning model recapitulates the main effects in our data.

### Supplementary Figures

#### Association of self-related learning rates and Psychopathology scores separately for both groups

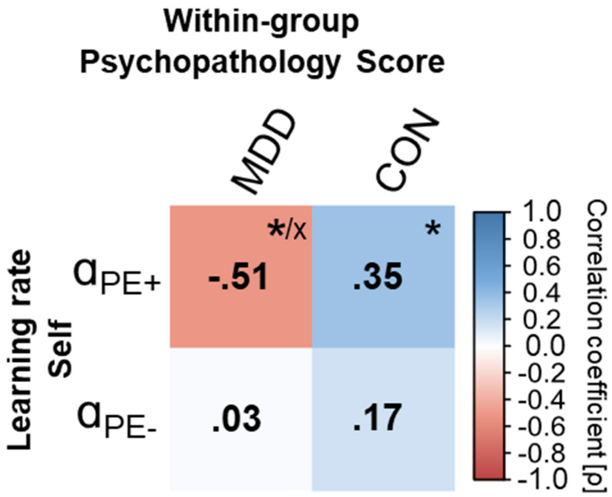


*Supplementary Figure 1*. Spearman correlation between self-related learning rates for positive ($\alpha_{PE+}$) and negative prediction errors ($\alpha_{PE-}$) with Psychopathology score for MDD and CON group. MDD = Major depressive disorder, CON = Control, * $p < .05$, x FDR corrected.

**Association between learning rates and symptom scores separately for both groups**

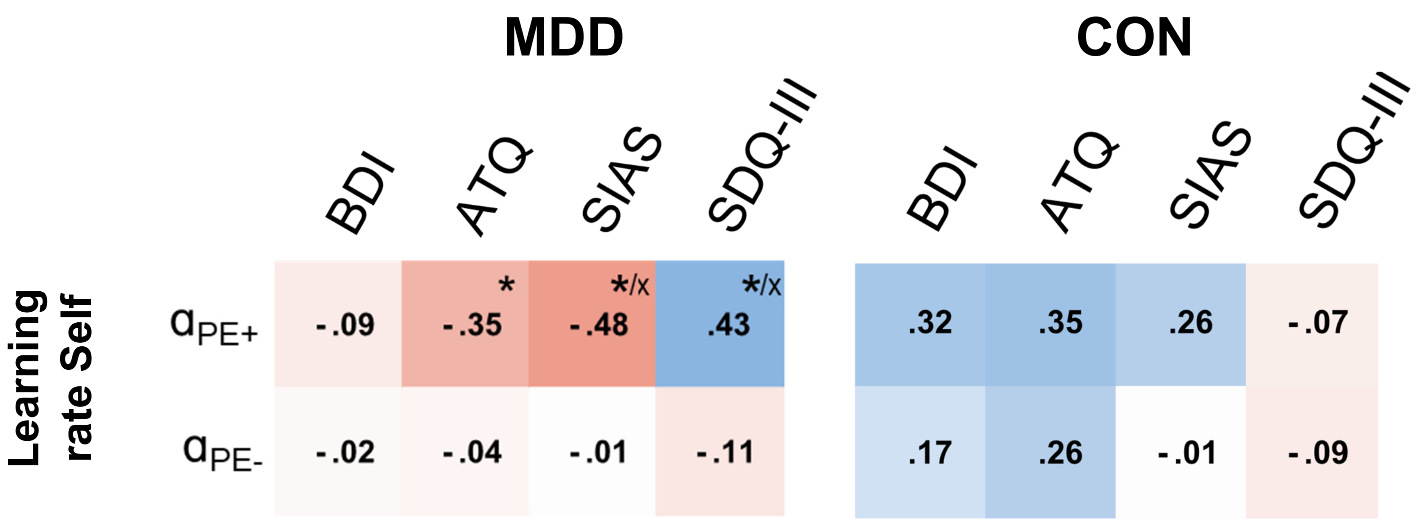


*Supplementary Figure 2.* Spearman correlation between the self-related learning rates for positive ($\alpha_{PE+}$) and negative prediction errors ($\alpha_{PE-}$) with the separate elements of the Psychopathology score: Beck's depression inventory (BDI-V[3]), Automatic Thought Questionnaire (ATQ[4]), Social Interaction Anxiety scale (SIAS[5]), Self-Description Questionnaire-III (SDQ-III subscale scores[6]). MDD = Major depressive disorder, CON = Control, * $p < .05$, x FDR corrected.

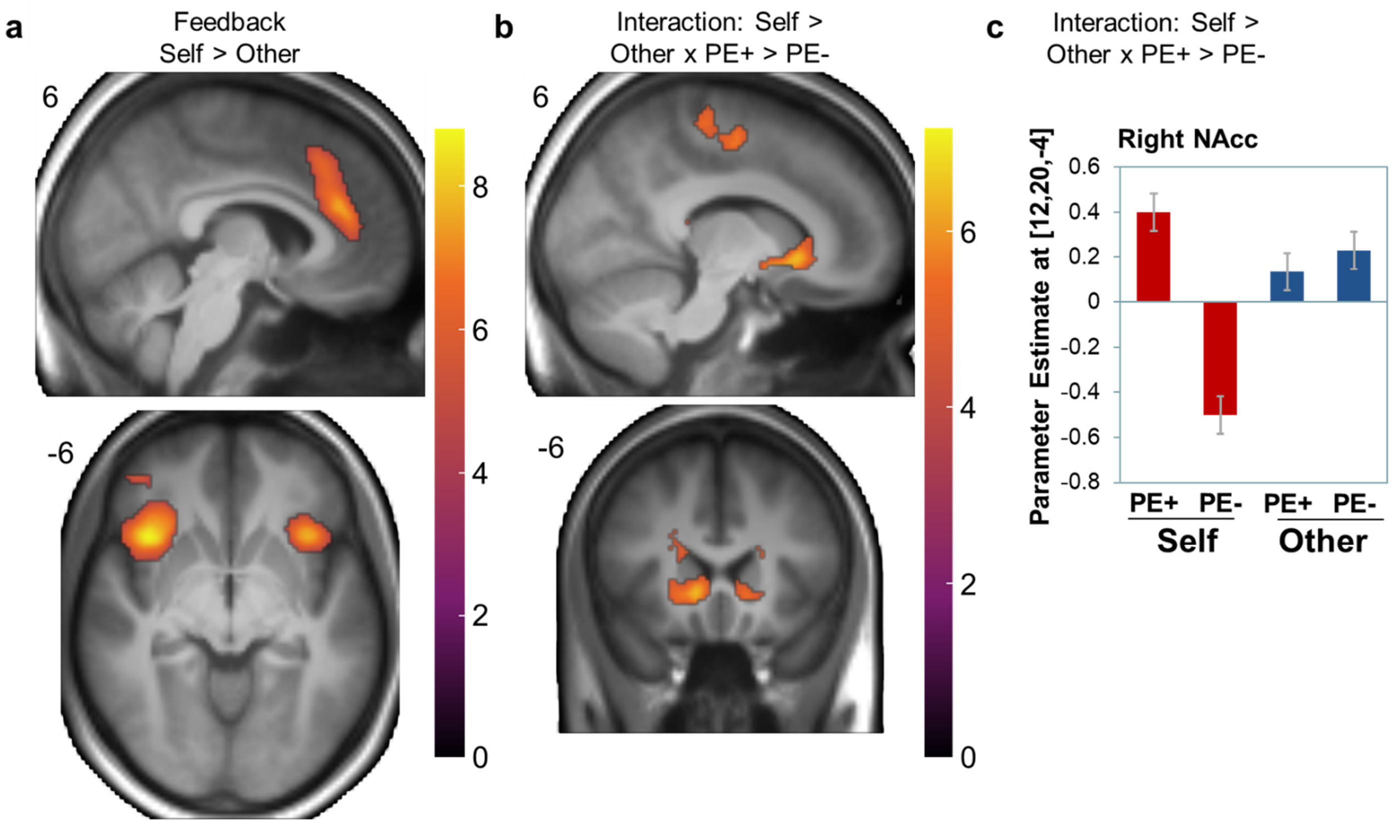


*Supplementary Figure 3.* Neural activations associated with prediction error processing (categorical PE effect) in the feedback phase. Replication of a previous study[7]. a) Self-related feedback vs. other-related feedback was associated with increased activation of the bilateral insula cortex, frontal orbital cortex, cingulate gyrus, supramarginal gyrus ($p < .05$, FWE corrected at peak level for the whole brain). b) The interaction of Agent and Prediction Error Valence ([Self PE+ > Self PE-] **>** [Other PE+ > Other PE-]) resulted in activation of the bilateral VS, precentral/ postcentral gyrus, left hippocampus ($p < .05$, FWE corrected at peak level for the whole brain). Anatomical labels were derived from the SPM Anatomy Toolbox Version 3.0. c) Means and standard errors of parameter estimates corresponding to the BOLD response to positive and negative PEs in the right VS: More activity in VS for positive relative to negative PEs only when seeing self-related feedback, but not when observing others.

Neural activation associated with negative compared to positive prediction error processing – whole brain analysis

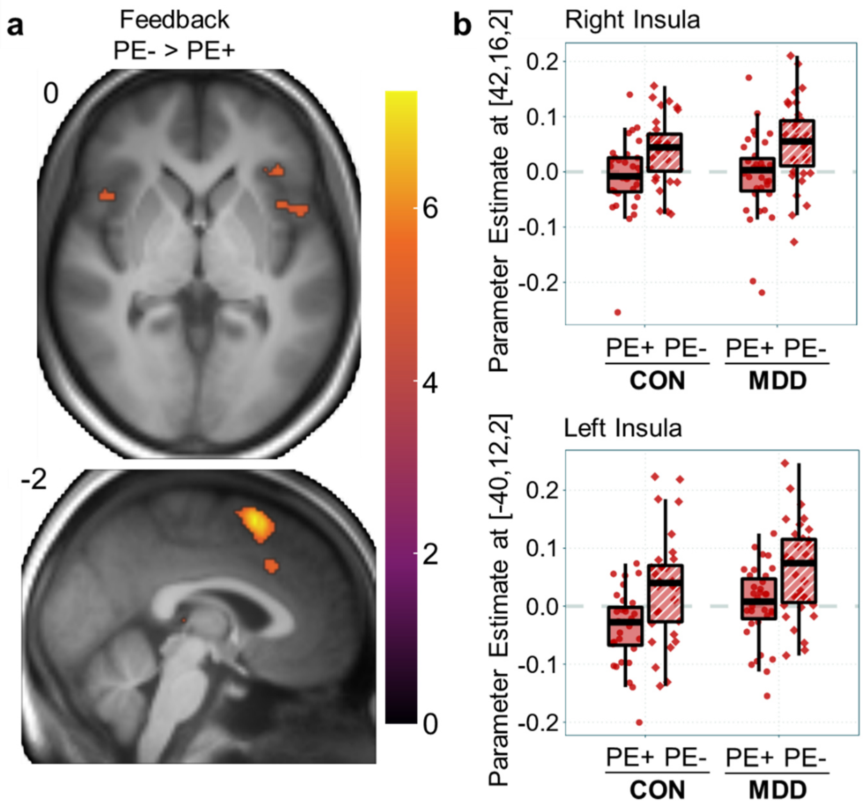


*Supplementary Figure 4.* Neural activation associated with negative compared to positive prediction error processing (continuous PE-Valence effect). a) Processing of self-related negative prediction errors (PE-) relative to positive prediction errors (PE+) was associated with increased activation of the superior frontal gyrus, paracingulate/ cingulate gyrus, frontal orbital cortex, operculum/ insula cortex ($p < .05$, FWE corrected at peak level for the whole brain). Anatomical labels were derived from the SPM Anatomy Toolbox Version 3.0. b) Parameter estimates for positive and negative prediction error processing at peak level within the insula cluster.

**Association of affective experience and brain activity for positive and negative prediction errors within the clinical sample**

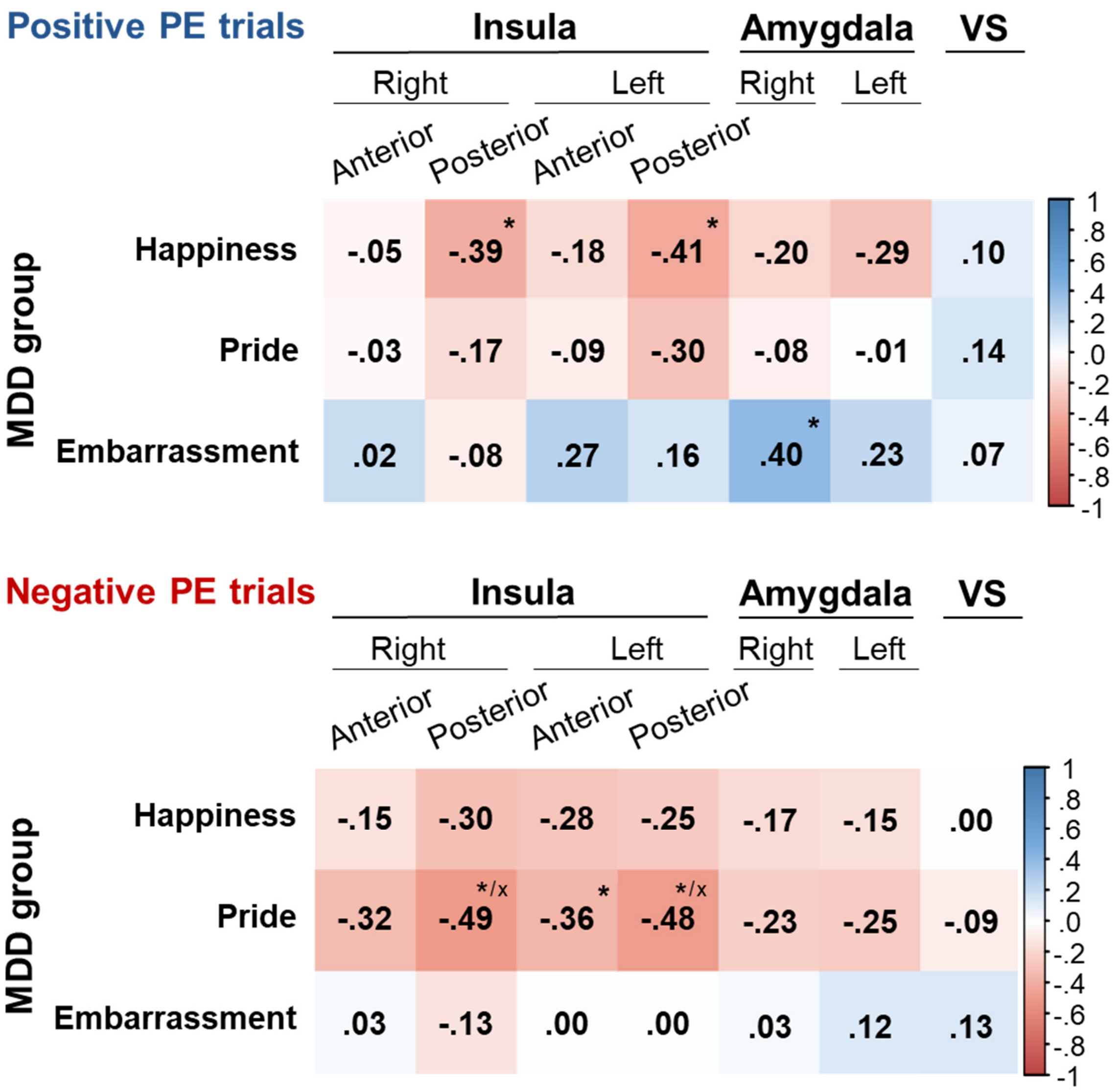


*Supplementary Figure 5.* Spearman correlation between the affect ratings and brain activity within ROIs in response to positive and negative prediction errors (categorical PE effect) in the clinical sample. * $p < .05$, x FDR corrected.

**Association of learning rates and brain activity for positive and negative prediction errors in the clinical sample**

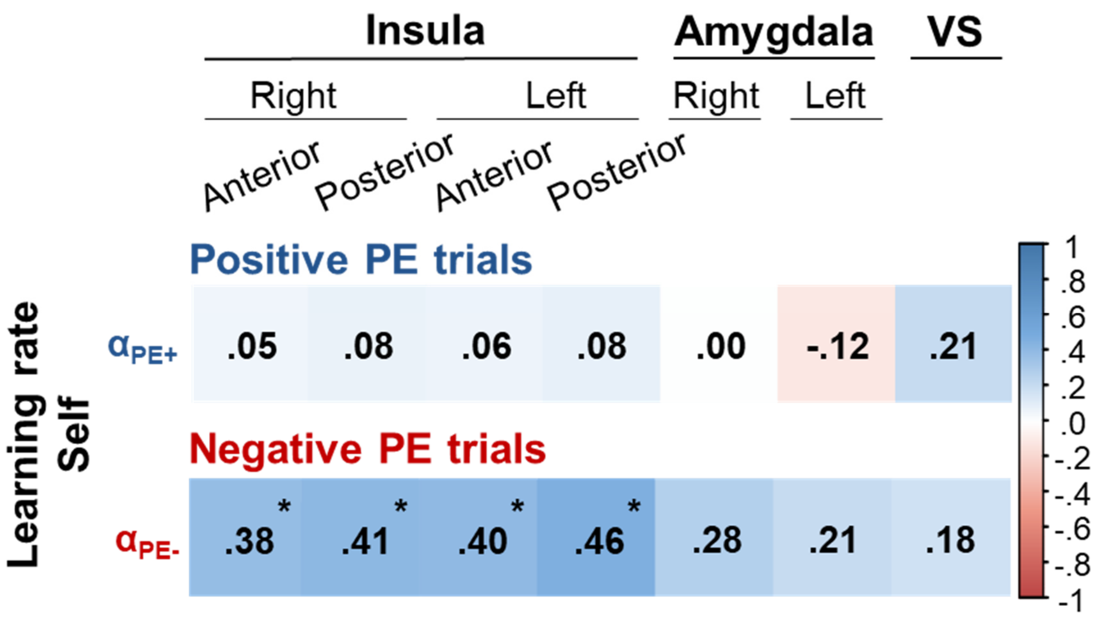


*Supplementary Figure 6.* Spearman correlation between learning rates for self-related positive (αPE+) and negative prediction errors (αPE-) and brain activity within ROIs in response to PE+ and PE- (categorical PE effect) in the clinical sample. * $p < .05$.

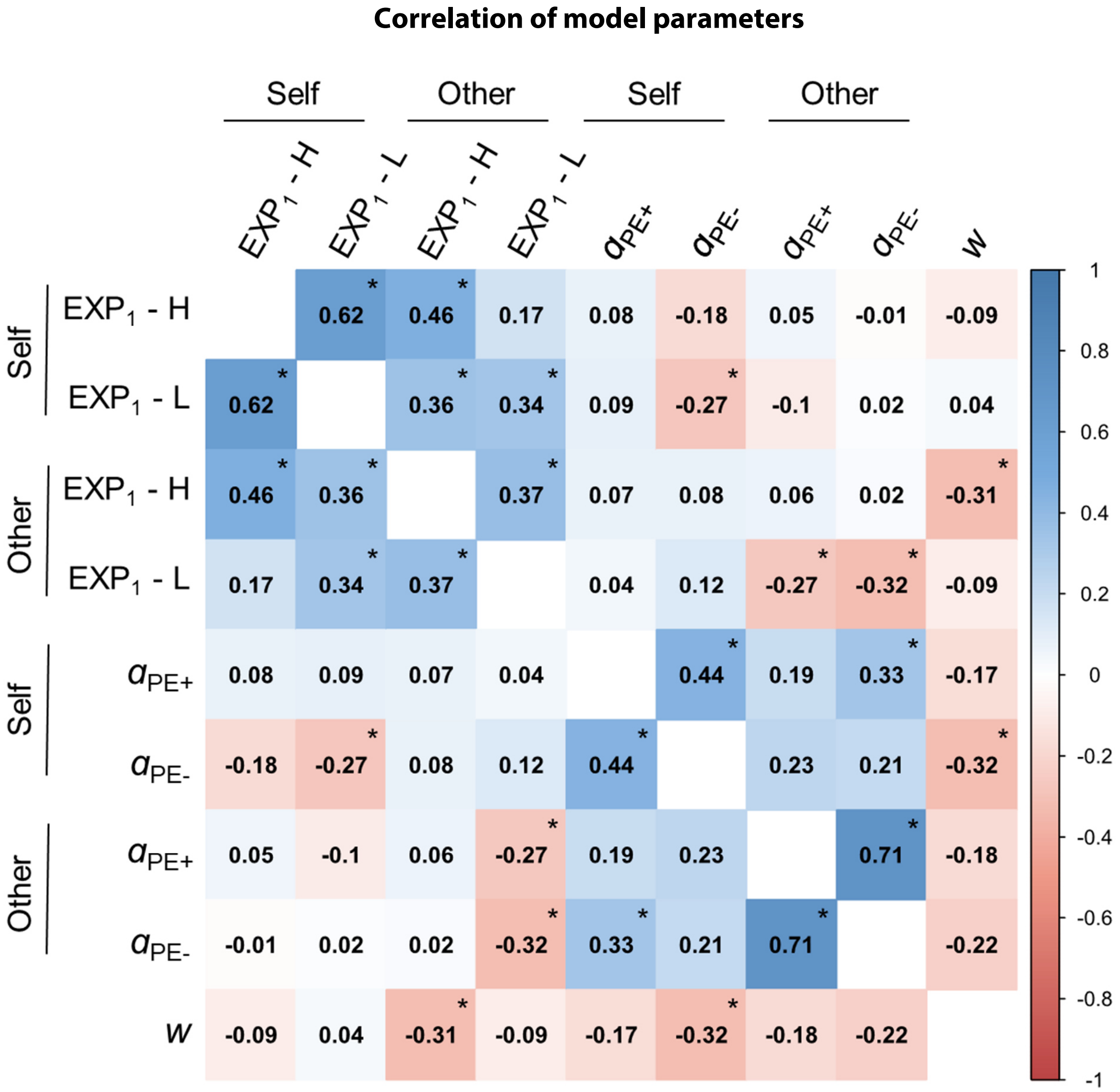


*Supplementary Figure 7.* Spearman correlation of model parameters for the winning model. $EXP_1$ = estimated first expected performance for the two ability conditions (H = high, L = low), separately for Self and Other. α = learning rates for positive (αPE+) and negative prediction errors (αPE-), separately for Self and Other. w = weighting factor. * $p < .05$.

**Supplementary Tables**

*Supplementary Table 1a.* Sample characteristics - Group comparison of age and psychometric tests

| | CON | | MDD | | *t*-test | | |
|---|---|---|---|---|---|---|---|
| | *M* | *SD* | *M* | *SD* | *t* | *p* | *d* |
| Age | 34.25 | 10.23 | 34.23 | 10.54 | 0.01 | 0.995 | 0.00 |
| BDI sum | 13.88 | 8.00 | 53.91 | 12.00 | -16.20 | <.001 | -3.88 |
| ATQ mean | 1.87 | 0.29 | 3.49 | 0.55 | -15.22 | <.001 | -3.63 |
| SIAS mean | 0.90 | 0.41 | 1.98 | 0.73 | -7.55 | <.001 | -1.80 |
| SDQ-III mean | 6.66 | 0.56 | 4.50 | 1.44 | 8.23 | < .001 | 1.96 |

*Note.* Group comparison of sample characteristics using a two-sample *t*-test. Beck's depression inventory (BDI-V[3], items on a 6-point Likert scale from zero to five), Automatic Thought Questionnaire (ATQ[4], items on a 5-point Likert scale from one to five), Social Interaction Anxiety scale (SIAS[5], items on a 5-point Likert scale from zero to four). Self-esteem measured with the Self-Description Questionnaire-III (SDQ-III[6], subscale score). *M* = mean, *SD* = standard deviation, *t* = *t*-value with Welch correction, *d* = Cohen's *d* with Hedges correction. MDD = Major depressive disorder (*n* = 35), CON = control group (*n* = 32).

*Supplementary Table 1b.* Sample characteristics - Group comparison of prior beliefs about global and specific estimation abilities

| | CON | | MDD | | *t*-test | | |
|---|---|---|---|---|---|---|---|
| | *M* | *SD* | *M* | *SD* | *t* | *p* | *d* |
| Global estimation ability - self | 5.09 | 1.00 | 4.49 | 1.36 | 2.10 | .040 | 0.50 |
| Specific estimation ability - self | 41.14 | 18.41 | 36.51 | 17.81 | 1.04 | .301 | 0.25 |
| Confidence of sp. estimation ability - self | 26.44 | 12.23 | 25.03 | 12.13 | 0.47 | .638 | 0.11 |
| Specific estimation ability - other | 49.12 | 14.64 | 46.21 | 14.38 | 0.82 | .415 | 0.20 |
| Confidence of sp. estimation ability - other | 34.79 | 14.97 | 25.46 | 13.99 | 2.63 | .011 | 0.64 |

*Note.* Group comparison of self-beliefs and beliefs about the other person rated in a pre-survey prior to the main task. Prior rating of global estimation ability (one item): 'I'm good at estimating,' (8-point Likert scale); mean of specific estimation ability ratings for the two self-related estimation categories (one item per category): 'How good are you at estimating [e.g., the weight of animals]?'; mean of two corresponding confidence ratings: 'How sure are you that your assessment of your ability is correct?'; mean of specific estimation ability ratings for the two other-related estimation categories: 'How good is the other person at estimating [e.g., the height of buildings]?'; mean of two corresponding other-related confidence ratings: 'How sure are you that your assessment of the other person's ability is correct?'. Specific estimation ability ratings on a scale from zero to 100% in comparison to a reference group; confidence ratings on a confidence interval scale from '+/- 50% (very unconfident)' to '+/- 0% (very confident)'. *M* = mean, *SD* = standard deviation, *t* = *t*-value with Welch correction, *d* = Cohen's *d* with Hedges correction. MDD = Major depressive disorder (*n* = 35), CON = control group (*n* = 32).

*Supplementary Table 1c.* Sample characteristics - Group comparison of setting characteristics regarding the estimation tasks and the other person prior to the task

| | CON | | MDD | | *t*-test | | |
|---|---|---|---|---|---|---|---|
| | *M* | *SD* | *M* | *SD* | *t* | *p* | *d* |
| Category-specific estimation experience | 2.44 | 1.32 | 3.01 | 1.27 | -1.82 | .073 | -0.44 |
| Importance of estimation ability | 2.80 | 1.39 | 3.17 | 1.40 | -1.10 | .277 | -0.26 |
| Familiarity of the other person | 1.09 | 0.3 | 1.20 | 0.41 | -1.23 | .223 | -0.29 |
| Likeability of other person | 5.16 | 1.14 | 5.20 | 1.23 | -0.15 | .880 | -0.04 |
| Estimated own likeability rated by other | 4.69 | 1.06 | 4.09 | 1.60 | 1.83 | .072 | 0.43 |

*Note.* Group differences in previous experience, importance of estimating, and attitude towards the other person. All characteristics were rated with one item each on a 7-point Likert scale in a pre-survey prior to the main task. Item wordings: 1. 'How much experience do you have with estimating [e.g., the weight of animals]?', 2. 'How important is it to you to be good at estimating [e.g., the weight of animals]?', 3. 'How well do you know the other person?', 4. 'I like the other person.', 5. 'I think the other person finds me likable.' *M* = mean, *SD* = standard deviation, *t* = *t*-value with Welch correction, *d* = Cohen's *d* with Hedges correction. MDD = Major depressive disorder (*n* = 35), CON = control group (*n* = 32).

*Supplementary Table 2.* Model-based analysis of belief updating behavior (learning rates) in individuals with and without depression

| | *Estimate* | *SE* | 95% CI | | *df* | *t* | *p* |
|---|---|---|---|---|---|---|---|
| | | | lower | upper | | | |
| (Intercept) | 0.22 | 0.03 | 0.17 | 0.27 | 195 | 8.83 | < .001 |
| PE-Valence [PE+] | 0.04 | 0.03 | -0.02 | 0.10 | 195 | 1.29 | .199 |
| Agent [Self] | 0.07 | 0.03 | 0.01 | 0.13 | 195 | 2.35 | .020 |
| Group | -0.01 | 0.04 | -0.08 | 0.06 | 65 | -0.30 | .764 |
| PE-Valence x Agent | -0.14 | 0.04 | -0.23 | -0.06 | 195 | -3.29 | .001 |
| PE-Valence x Group | -0.01 | 0.04 | -0.09 | 0.07 | 195 | -0.23 | .818 |
| Agent x Group | 0.03 | 0.04 | -0.06 | 0.11 | 195 | 0.62 | .533 |
| PE-Valence x Agent x Group | 0.01 | 0.06 | -0.11 | 0.13 | 195 | 0.18 | .861 |

*Note.* Prediction Error Valence (PE-Valence, [PE+, PE-]) x Agent [Self, Other] x Group [MDD, CON] linear mixed-effects model fit by maximum likelihood. Dependent variable: learning parameters of the winning model. Beta-estimate, *SE* = standard error, *CI* = confidence interval, *df* = degrees of freedom, *t* = *t*-value, MDD = Major depressive disorder ($n$ = 35), CON = control group ($n$ = 32).

*Supplementary Table 3.* Model Space

| Assumptions about learning | Model | Learning Parameters |
|---|---|---|
| **Self = Other** | | |
| No learning bias | Unity Model (M1) | ***a*** |
| Prediction Error: Positive ≠ Negative | Valence Model (M2) | ***a***PE+, ***a***PE- |
| Ability Context: High ≠ Low | Ability Model (M3) | ***a***High, ***a***Low |
| **Self ≠ Other** | | |
| No learning bias | Unity Model (M4) | ***a***Self<br>***a***Other |
| Prediction Error: Positive ≠ Negative | Valence Model (M5) | ***a***Self/PE+, ***a***Self/PE-<br>***a***Other/PE+, ***a***Other/PE- |
| Prediction Error: Positive ≠ Negative & Decay for extreme feedback values | Weighted Valence Model (M6) | ***a***Self/PE+, ***a***Self/PE-<br>***a***Other/PE+, ***a***Other/PE-<br>***w*** |
| Ability Context: High ≠ Low | Ability Model (M7) | ***a***Self/High, ***a***Self/Low<br>***a***Other/High, ***a***Other/Low |
| **No learning** | Mean Model (M0) | |

*Note:* Four starting values (estimated $EXP_1$) for the four feedback conditions (Agent [Self vs. Other] x Ability [High Ability vs. Low Ability] were additionally estimated for all models.

*Supplementary Table 4.* Model comparison

| *Model* | | *PSIS-LOO* | *LOO-SE* | *LOO-Diff* | *% of $\hat{k}$ > 0.7* | *Num. param.* |
|---|---|---|---|---|---|---|
| **Whole sample** | | | | | | |
| | Mean Model (M0) | -2441.5 | 216.15 | -1176.0 | 0.04 | 4 |
| Self = Other | Unity Model (M1) | -1722.6 | 216.37 | -457.1 | 0.62 | 5 |
| | Valence Model (M2) | -1580.0 | 210.76 | -314.5 | 0.34 | 6 |
| | Ability Mode (M3) | -1639.2 | 213.23 | -373.7 | 0.71 | 6 |
| Self ≠ Other | Unity Model (M4) | -1644.7 | 209.58 | -379.2 | 0.49 | 6 |
| | Valence Model (M5) | -1356.1 | 219.24 | -90.6 | 0.39 | 8 |
| | Weighted Valence Model (M6) | -1265.5 | 229.54 | - | 1.10 | 9 |
| | Ability Model (M7) | -1539.0 | 208.47 | -273.4 | 1.32 | 8 |
| **MDD** | | | | | | |
| | Mean Model (M0) | -1404.3 | 156.32 | -603.8 | 0.02 | 4 |
| Self = Other | Unity Model (M1) | -1089.8 | 182.84 | -289.3 | 0.49 | 5 |
| | Valence Model (M2) | -1000.3 | 176.65 | -199.8 | 0.19 | 6 |
| | Ability Model (M3) | -1039.4 | 179.52 | -238.9 | 0.58 | 6 |
| Self ≠ Other | Unity Model (M4) | -1022.5 | 175.15 | -222.0 | 0.22 | 6 |
| | Valence Model (M5) | -835.4 | 182.05 | -34.9 | 0.17 | 8 |
| | Weighted Valence Model (M6) | -800.5 | 191.68 | - | 0.58 | 9 |
| | Ability Model (M7) | -961.9 | 173.61 | -161.4 | 0.76 | 8 |
| **CON** | | | | | | |
| | Mean Model (M0) | -1037.2 | 148.25 | -572.1 | 0.02 | 4 |
| Self = Other | Unity Model (M1) | -632.8 | 108.87 | -167.8 | 0.13 | 5 |
| | Valence Model (M2) | -579.8 | 109.48 | -114.7 | 0.15 | 6 |
| | Ability Model (M3) | -599.9 | 108.84 | -134.8 | 0.13 | 6 |
| Self ≠ Other | Unity Model (M4) | -622.2 | 110.69 | -157.2 | 0.26 | 6 |
| | Valence Model (M5) | -520.8 | 121.02 | -55.7 | 0.22 | 8 |
| | Weighted Valence Model (M6) | -465.0 | 124.64 | - | 0.52 | 9 |
| | Ability Model (M7) | -577.0 | 111.48 | -112.0 | 0.56 | 8 |

*Note.* LOO = sum PSIS-LOO, approximate leave-one-out cross-validation (LOO) using Pareto-smoothed importance sampling (PSIS); LOO-SE = Standard error of PSIS-LOO; LOO-Diff (SE-Diff) = Difference in expected predictive accuracy (PSIS-LOO) for all models from the model with the highest PSIS-LOO (weighted Valence Model) and standard errors of differences; percentage of $\hat{k}$ - estimated shape parameters of the generalized Pareto distribution - exceeding 0.7 (all according to Vehtari et al.); Num. param. = number of estimated parameters in the model; Self = Other: same learning rates for Self and Other, Self ≠ Other: separate learning rates for Self and Other; MDD = Major depressive disorder (*n* = 35), CON = control group (*n* = 32).

*Supplementary Table 5.* Model-agnostic analysis of belief-updating behavior in individuals with and without depression

| | *Estimate* | *SE* | 95% CI | | *df* | *t* | *p* |
|---|---|---|---|---|---|---|---|
| | | | lower | upper | | | |
| (Intercept) | 51.94 | 1.35 | 49.31 | 54.58 | 5279 | 38.61 | < .001 |
| Ability [High] | 5.49 | 1.25 | 3.04 | 7.94 | 5279 | 4.39 | < .001 |
| Agent [Self] | -3.62 | 1.25 | -6.06 | -1.17 | 5279 | -2.89 | .004 |
| Trial | -0.56 | 0.07 | -0.70 | -0.41 | 5279 | -7.54 | < .001 |
| **Group** [MDD] | -1.19 | 1.86 | -4.90 | 2.53 | 65 | -0.64 | .526 |
| Ability x Agent | -0.81 | 1.77 | -4.27 | 2.65 | 5279 | -0.46 | .649 |
| Ability x Trial | 1.08 | 0.10 | 0.88 | 1.28 | 5279 | 10.34 | < .001 |
| Ability x **Group** | -0.86 | 1.73 | -4.24 | 2.53 | 5279 | -0.49 | .621 |
| Agent x Trial | -0.27 | 0.10 | -0.47 | -0.07 | 5279 | -2.59 | .010 |
| Agent x **Group** | -1.90 | 1.73 | -5.29 | 1.48 | 5279 | -1.10 | .271 |
| Trial x **Group** | 0.09 | 0.10 | -0.11 | 0.29 | 5279 | 0.85 | .394 |
| Ability x Agent x Trial | 0.07 | 0.15 | -0.22 | 0.36 | 5279 | 0.45 | .649 |
| Ability x Agent x **Group** | 0.36 | 2.45 | -4.42 | 5.15 | 5279 | 0.15 | .882 |
| Ability x Trial x **Group** | -0.08 | 0.14 | -0.36 | 0.21 | 5279 | -0.54 | .593 |
| Agent x Trial x **Group** | -0.03 | 0.14 | -0.32 | 0.25 | 5279 | -0.23 | .819 |
| Ability x Agent x Trial x **Group** | 0.08 | 0.20 | -0.32 | 0.48 | 5279 | 0.39 | .697 |

*Note.* Trial (continuous, 1-20) x Agent [Self, Other] x Ability condition [High, Low] x Group [MDD, CON] linear mixed-effects model fit by maximum likelihood. Dependent variable: trial-by-trial ratings of expected performance. Beta-estimate, *SE* = standard error, *CI* = confidence interval, *df* = degrees of freedom, *t* = t-value, MDD = Major depressive disorder (*n* = 35), CON = control group (*n* = 32).

*Supplementary Table 6.* Group comparison of positive and negative prediction error tracking within regions of interest

| Covariates/ Regions of interest | Side | Cluster Size | MNI Coordinates | | | *T* | *p* |
|---|---|---|---|---|---|---|---|
| | | | x | y | z | | |
| **Interaction Group x PE-Valence** | | | | | | | |
| MDD (PE-), CON (PE+) > MDD (PE+), CON (PE-) | | | | | | | |
| Anterior insula - right | R | 85 | 38 | -12 | 2 | 3.35 | .045 |
| Anterior insula - left | L | 32 | -38 | 14 | -10 | 2.40 | .309 |
| Posterior insula - right | R | 175 | 36 | -18 | 4 | 3.17 | .030 |
| Posterior insula - left | L | 3 | -30 | -26 | 18 | 2.03 | .328 |
| Amygdala - right | R | 58 | 34 | 2 | -16 | 2.27 | .236 |
| Amygdala - left | L | 39 | -26 | -6 | -18 | 2.75 | .086 |
| Ventral Striatum (functional PE ROI) | No suprathreshold clusters | | | | | | |
| **Positive prediction error - CON > MDD** | | | | | | | |
| All ROIs | No suprathreshold clusters | | | | | | |
| **Negative prediction error - MDD > CON** | | | | | | | |
| Anterior insula - right | R | 627 | 36 | 16 | -8 | 3.11 | .059 |
| Anterior insula - left | L | 138 | -40 | 16 | -10 | 2.67 | .151 |
| Posterior insula - right | R | 272 | 40 | -18 | 12 | 3.30 | .014 |
| Posterior insula - left | L | 33 | -36 | -22 | 14 | 2.39 | .166 |
| Amygdala - right | R | 241 | 34 | 0 | -12 | 2.48 | .129 |
| Amygdala - left | L | 107 | -24 | -2 | -18 | 2.58 | .108 |
| Ventral Striatum (functional PE ROI) | No suprathreshold clusters | | | | | | |

*Note.* Group comparison of prediction error tracking for positive (PE+) and negative (PE-) prediction errors using two-sample *t*-tests and Group x PE-Valence interaction using a flexible factorial design. The *p*-values are FWE corrected within ROIs at peak level. All three tests had no suprathreshold clusters on the whole brain level (FWE-corrected). *R* = right, *L* = left. *T* = *t*-value. MDD = Major depressive disorder ($n$ = 35), CON = control group ($n$ = 32).

*Supplementary Table 7.* Baseline activations associated with the tracking of positive and negative prediction errors

| Contrasts/ Brain regions | Side | Cluster Size | MNI Coordinates x | y | z | *T* | *p* |
|---|---|---|---|---|---|---|---|
| **Positive prediction error** | | | | | | | |
| Angular Gyrus/ Lateral Occipital Cortex, superior division | R | 69 | -56 | -60 | 24 | 5.40 | .012 |
| Angular Gyrus/ Supramarginal Gyrus, posterior division | | | -46 | -50 | 24 | 5.20 | .022 |
| Angular Gyrus/ Lateral Occipital Cortex, superior division | L | 8 | 52 | -54 | 24 | 5.15 | .026 |
| **Negative prediction error** | | | | | | | |
| Superior Frontal Gyrus/ Juxtapositional Lobule Cortex | R/ L | 245 | -2 | 12 | 60 | 7.29 | <.001 |
| Inferior Frontal Gyrus, pars triangularis/ pars opercularis | L | 145 | 48 | 28 | 4 | 6.34 | <.001 |
| | | | 50 | 20 | 2 | 5.45 | .009 |
| Frontal Orbital Cortex/ Frontal Operculum Cortex | | | 40 | 28 | -4 | 5.30 | .014 |
| Superior Frontal Gyrus/ Paracingulate Gyrus | | 65 | -6 | 52 | 22 | 5.59 | .005 |
| Paracingulate Gyrus/ Superior Frontal Gyrus | | | 8 | 50 | 22 | 5.48 | .008 |
| Frontal Orbital Cortex/ Frontal Operculum Cortex | | 19 | -38 | 26 | -6 | 5.23 | .017 |

*Note*. Baseline activations of prediction error tracking. Positive and negative prediction error refer to the unsigned prediction error values as two parametric modulators for the feedback phase of the Self condition. The *p*-values are FWE-corrected for the whole brain at peak level. Whole sample: $n = 67$. *R* = right, *L* = left. *T* = *t*-value. Anatomical labels were derived from the SPM Anatomy Toolbox Version 3.0.

*Supplementary Table 8.* Emotions - group comparison

| | CON | | MDD | | *t*-test | | |
|---|---|---|---|---|---|---|---|
| | *M* | *SD* | *M* | *SD* | *t* | *p* | *d* |
| Happiness | 58.28 | 12.04 | 42.98 | 21.95 | 3.58 | < .001 | 0.85 |
| Arousal | 30.41 | 20.89 | 41.70 | 23.91 | -2.06 | .043 | -0.50 |
| Pride | 45.06 | 19.66 | 34.73 | 22.62 | 2.00 | .050 | 0.48 |
| Embarrassment | 20.34 | 18.61 | 23.48 | 23.91 | -0.60 | .549 | -0.14 |
| Tiredness | 34.47 | 22.41 | 44.56 | 26.32 | -1.69 | .095 | -0.41 |

*Note.* Group comparison of emotion ratings during task performance. Two ratings per emotion on a continuous scale from zero to 100 following the presentation of self-related feedback trials. *M* = mean, *SD* = standard deviation, *d* = Cohen's *d* with Hedges correction. MDD = Major depressive disorder (*n* = 35), CON = control group (*n* = 32).
